# VPS35 and α-Synuclein Fail to Interact to Modulate Neurodegeneration in Rodent Models of Parkinson’s Disease

**DOI:** 10.1101/2022.12.07.519521

**Authors:** Xi Chen, Elpida Tsika, Nathan Levine, Darren J. Moore

## Abstract

Mutations in the *vacuolar protein sorting 35 ortholog* (*VPS35*) gene cause late-onset, autosomal dominant Parkinson’s disease (PD), with a single missense mutation (Asp620Asn, D620N) known to segregate with disease in families with PD. The *VPS35* gene encodes a core component of the retromer complex, involved in the endosomal sorting and recycling of transmembrane cargo proteins. *VPS35*-linked PD is clinically indistinguishable from sporadic PD, although it is not yet known whether *VPS35*-PD brains exhibit α-synuclein-positive brainstem Lewy pathology that is characteristic of sporadic cases. Prior studies have suggested a functional interaction between VPS35 and the PD-linked gene product α-synuclein in lower organisms, where *VPS35* deletion enhances α-synuclein-induced toxicity. In mice, VPS35 overexpression is reported to rescue hippocampal neuronal loss in human α-synuclein transgenic mice, potentially suggesting a retromer deficiency in these mice. Here, we employ multiple well-established genetic rodent models to explore a functional or pathological interaction between VPS35 and α-synuclein *in vivo*. We find that endogenous α-synuclein is dispensable for nigrostriatal pathway dopaminergic neurodegeneration induced by the viral-mediated delivery of human D620N VPS35 in mice, suggesting that α-synuclein does not operate downstream of VPS35. We next evaluated retromer levels in affected brain regions from human A53T-α-synuclein transgenic mice, but find normal levels of the core subunits VPS35, VPS26 or VPS29. We further find that heterozygous *VPS35* deletion fails to alter the lethal neurodegenerative phenotype of these A53T-α-synuclein transgenic mice, suggesting the absence of retromer deficiency in this PD model. Finally, we explored the neuroprotective capacity of increasing VPS35 expression in a viral-based human wild-type α-synuclein rat model of PD. However, we find that the overexpression of wild-type VPS35 is not sufficient for protection against α-synuclein-induced nigral dopaminergic neurodegeneration, α-synuclein pathology and reactive gliosis. Collectively, our data suggest a limited interaction of VPS35 and α-synuclein in neurodegenerative models of PD, and do not provide support for their interaction within a common pathophysiological pathway.

## Introduction

Parkinson’s disease (PD) is a complex neurodegenerative movement disorder that typically occurs in a sporadic manner, yet 5-10% of cases are inherited and monogenic (1-5). Among the familial forms of PD, mutations in the *Vacuolar Protein Sorting 35* (*VPS35*) gene cause late-onset, autosomal dominant PD (6-8). A single heterozygous mutation, Asp620Asn (D620N), has been identified to unambiguously segregate with disease in multiple families with PD and is the most frequent cause of *VPS35*-linked disease (9). Subjects with PD harboring *VPS35* mutations exhibit a clinical spectrum and neuroimaging findings indistinguishable from sporadic PD, although it is not yet clear whether brains from these subjects develop typical brainstem Lewy pathology (6-8, 10-12). Elucidating the mechanisms by which *VPS35* mutations cause PD are of central importance for defining common cellular pathways that drive neurodegeneration and for therapeutic development. Together with other PD-linked genes, the emergence of *VPS35* has highlighted a key role for the endolysosomal pathway in the pathophysiology of PD (13).

*VPS35* encodes a core component of the pentameric retromer complex that functions in the retrograde transport and recycling of transmembrane cargo proteins from endosomes to the *trans*-Golgi network or plasma membrane (14-17). Retromer is composed of a central cargo-selective complex that consists of VPS35 bound to VPS26 and VPS29, together with a sorting nexin dimer that plays a role in membrane binding and deformation (18). Much of what is known about the retromer, including the identification of its selective cargo, is derived from studies in yeast and mammalian cell lines, yet its role in brain cells remains poorly understood. How the PD-linked D620N mutation induces neuronal degeneration in PD is not yet clear (9, 19). D620N VPS35 may influence the endosomal sorting of specific cargo in certain cellular models (20-24), and is also known to impair the recruitment of the pentameric WASH complex to endosomes via a reduced interaction of VPS35 with the FAM21 subunit (21, 23). Reduced WASH binding to VPS35 may lead to altered vesicular sorting of the autophagy receptor ATG9A and impaired macroautophagy in mammalian cell lines (23). The D620N mutation in *VPS35* has also been linked to alterations in autophagy and mitochondrial morphology by regulating the chaperone-mediated autophagy receptor, LAMP2A (22), and the mitochondrial fusion/fission proteins, mitofusin-2 or Drp1 (25, 26), respectively.

Recent studies have suggested an intriguing relationship between VPS35 and α-synuclein (αSyn) (22, 25, 27, 28), a familial and risk gene for PD and the major component of Lewy bodies. In rodent brain, the heterozygous germline deletion of *VPS35*, or its conditional homozygous deletion in dopaminergic neurons, is reported to induce substantia nigra dopaminergic neuronal loss and promote the neuronal accumulation of αSyn (22, 25). Notably, the homozygous germline deletion of *VPS35* results in early embryonic lethality (29). *D620N VPS35* knockin mice, a more physiologically-relevant model of PD, also exhibit dopaminergic neurodegeneration yet it remains uncertain whether there is brain αSyn accumulation due to conflicting reports (30, 31). However, crossing the *D620N VPS35* knockin mice with human A53T-αSyn transgenic mice was not sufficient to alter the lethal neurodegenerative phenotype of these αSyn mice (30). Instead, these *VPS35* knockin mice exhibit the somatodendritic accumulation of abnormal tau protein throughout the brain (30). Furthermore, the viral-mediated expression of human D620N VPS35 in the nigrostriatal pathway of adult rats is sufficient to induce nigral dopaminergic neuronal loss yet in the absence of obvious αSyn accumulation or pathology (32). While there is some evidence suggesting that VPS35 can regulate αSyn accumulation (22, 25, 31), what is not yet clear is whether this is sufficient or necessary for driving neurodegeneration in these VPS35 models. There is also evidence for an opposite effect whereby pathological αSyn may drive a functional retromer deficiency. For example, *VPS35* deletion enhances toxic phenotypes induced by human αSyn expression in yeast, *C*.*elegans* and *Drosophila* models of PD (28, 33). The lentiviral-mediated overexpression of human wild-type (WT) VPS35 has been shown to rescue pyramidal neuronal loss, reactive astrogliosis and αSyn accumulation in the hippocampus of human WT-αSyn transgenic mice (28). These findings support the notion that αSyn can induce a retromer deficiency and that restoring one retromer subunit, VPS35, is sufficient to provide neuroprotection. Whether this reported protective effect of VPS35 is also relevant for vulnerable neuronal populations in PD requires further evaluation.

Here, we set out to better define the functional and pathological interaction between VPS35 and αSyn in the brain. We focus our studies on whether αSyn is required downstream of D620N VPS35 for mediating neurodegeneration in mice, and oppositely whether αSyn pathology can induce a downstream retromer deficiency using multiple rodent models. Our data demonstrate that endogenous αSyn is dispensable for D620N VPS35-induced dopaminergic neurodegeneration in mice, and we find no evidence for a physical or functional retromer deficiency contributing to human αSyn-induced neurodegeneration in rodents. Our study fails to provide support for a meaningful or robust interaction between VPS35 and αSyn in the brain of PD animal models, indicating that these proteins may operate in independent pathways in PD.

## Results

### _α_Syn does not interact with VPS35 or regulate retromer levels in human cells

To begin to explore the functional relationship between VPS35 and αSyn, we evaluated the interaction of both proteins by co-immunoprecipitation (co-IP) in HEK-293T cells transiently co-expressing V5-tagged VPS35 variants and untagged or Myc-tagged αSyn. IP of wild-type (WT) or PD-linked variants (P316S, D620N) of human VPS35 fails to detect a robust interaction with human WT αSyn by Western blot analysis (**Fig. 1A-B**), whereas VPS35 variants do interact with the endogenous retromer subunit VPS26 (**Fig. 1A**). Prior studies have suggested that αSyn overexpression induces a retromer deficiency (28). To evaluate this possibility, we transiently expressed untagged human αSyn variants in human SH-SY5Y cells. The overexpression of WT or PD-linked variants (A30P, E46K, A53T) of αSyn does not alter the steady-state levels of the endogenous retromer subunits VPS35, VPS26 or VPS29 by Western blot analysis (**Fig. 1C-D**). Previous studies have also suggested that VPS35 depletion induces the accumulation of αSyn (22, 25, 28). To test this idea, we generated SH-SY5Y cell clones stably expressing miR30-adapted short hairpin RNAs (shRNAs) targeting VPS35 or a non-silencing control, with or without the transient overexpression of untagged human WT αSyn. Two distinct shRNAs against VPS35 were successful in markedly reducing the levels of endogenous VPS35 protein that also induces a corresponding reduction in VPS26 and VPS29 (**Fig. 1E**). VPS35 depletion fails to increase the levels of endogenous αSyn protein in SH-SY5Y cells, but surprisingly causes a reduction in the levels of overexpressed WT αSyn (**Fig. 1E**). Similar experiments in these stable shRNA cells transiently expressing Myc-tagged human tau also reveals a reduction in tau protein levels induced by VPS35 depletion (**Fig. 1F**), most likely suggesting that VPS35-shRNA cells exhibit poor transfection efficiency relative to non-silencing shRNA cells. Taken together, our data initially fails to provide evidence for i) a biochemical interaction of VPS35 with αSyn, ii) an αSyn-induced retromer deficiency, and iii) VPS35 depletion inducing αSyn accumulation, in human cells.

**Figure 1:**
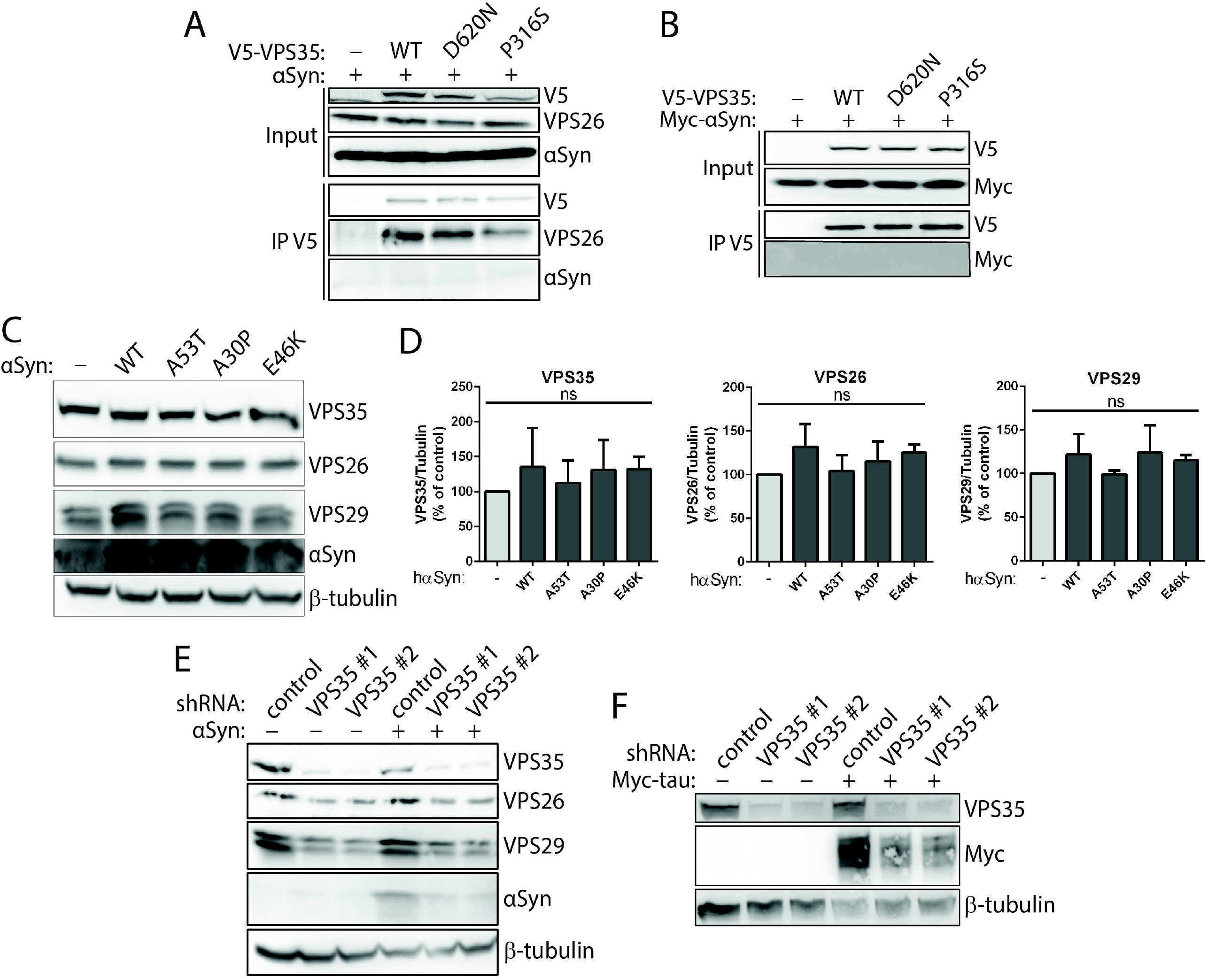
Lack of interaction of _α_Syn and VPS35 in human cells. **A-B**) Co-IP assays between V5-tagged human VPS35 (WT, D620N and P316S) and (**A**) untagged or (**B**) Myc-tagged human WT αSyn indicate a lack of interaction of VPS35 with αSyn. HEK-293T cell extracts co-expressing V5-tagged VPS35 (WT, D620N or P316S) and WT αSyn were subjected to IP with anti-V5 antibody, and IP and input fractions were probed for αSyn, Myc (αSyn), VPS26 and V5. VPS35 variants interact equivalently with endogenous VPS26 but not with αSyn. **C-D**) Overexpression of PD-linked αSyn variants does not influence the steady-state levels of endogenous retromer subunits. **C-D**) Western blot analysis of Triton-soluble extracts from SH-SY5Y cells transiently expressing untagged human αSyn variants (WT, A53T, A30P or E46K) or empty vector. Graphs indicate the levels of each retromer subunit, VPS35, VPS26 or VPS29, normalized to β-tubulin levels (mean ± SEM, *n* = 3 experiments). Data were analyzed by one-way ANOVA with Dunnett’s *post-hoc* analysis. *ns*, non-significant. **E**) Knockdown of endogenous VPS35 does not increase the levels of endogenous or overexpressed human WT αSyn. Western blot analysis of Triton-soluble extracts from SH-SY5Y cell clones stably expressing miR30-adapted shRNAs targeting VPS35 (#1 or #2) or a non-silencing control shRNA. VPS35 levels are markedly reduced in cells expressing VPS35-shRNAs together with corresponding reductions in VPS26 and VPS29, as expected, whereas overexpressed αSyn is also reduced. **F**) Western blot analysis of SH-SY5Y cells stably expressing VPS35-shRNAs reveal a similar reduction of overexpressed Myc-tagged human WT tau.

### _α_Syn is not required for dopaminergic neurodegeneration induced by the overexpression of human D620N VPS35 in mice

It is not yet known whether PD brains harboring *VPS35* mutations are associated with Lewy pathology (7), and *D620N VPS35* knockin mouse models do not appear to exhibit αSyn aggregation with advancing age (30). Since our experiments in cell lines did not reveal an interaction between VPS35 and αSyn, we turned to rodent models to first evaluate whether endogenous αSyn is functionally required for mediating the pathogenic actions of PD-linked D620N VPS35. We unilaterally delivered recombinant AAV2/6 vectors expressing V5-tagged human D620N VPS35 (or a control virus containing a stuffer sequence, MCS) to the substantia nigra of age-matched adult homozygous *SNCA* (αSyn) knockout (KO) or WT littermate mice. We have previously demonstrated that the viral-mediated delivery of D620N VPS35 induces a progressive ∼30% loss of nigral dopaminergic neurons in adult mice and rats (32, 34). At 12 weeks post-injection in mice, we observe robust expression of D620N VPS35 limited to the injected ipsilateral substantia nigra pars compacta of WT and *SNCA* KO mice that substantially co-localizes with the dopaminergic neuronal marker, tyrosine hydroxylase (TH) (**Fig. 2A**). To evaluate the impact of αSyn deletion on neurodegeneration, unbiased stereology was used to count the number of TH-positive dopaminergic neurons and total Nissl-positive neurons in the ipsilateral (injected) compared to the contralateral (non-injected) nigra. D620N VPS35 expression induces a marked loss of nigral dopaminergic neurons in WT mice that is not significantly altered in KO mice, relative to the negligible effects of a control virus (**Fig. 2B-C**). The loss of TH-positive neurons in all conditions is paralleled by a comparable loss of total Nissl-positive neurons (**Fig. 2C**), confirming dopaminergic neuronal degeneration rather than a loss of TH phenotype. D620N VPS35 expression also induces a marked yet non-significant loss of TH-positive dopaminergic nerve terminals in the striatum, yet with no difference between WT and KO mice (**Fig. 2D-E**). Our data suggest that endogenous αSyn is not critically required for dopaminergic neurodegeneration induced by the viral-mediated expression of D620N VPS35 in the mouse nigrostriatal pathway.

**Figure 2:**
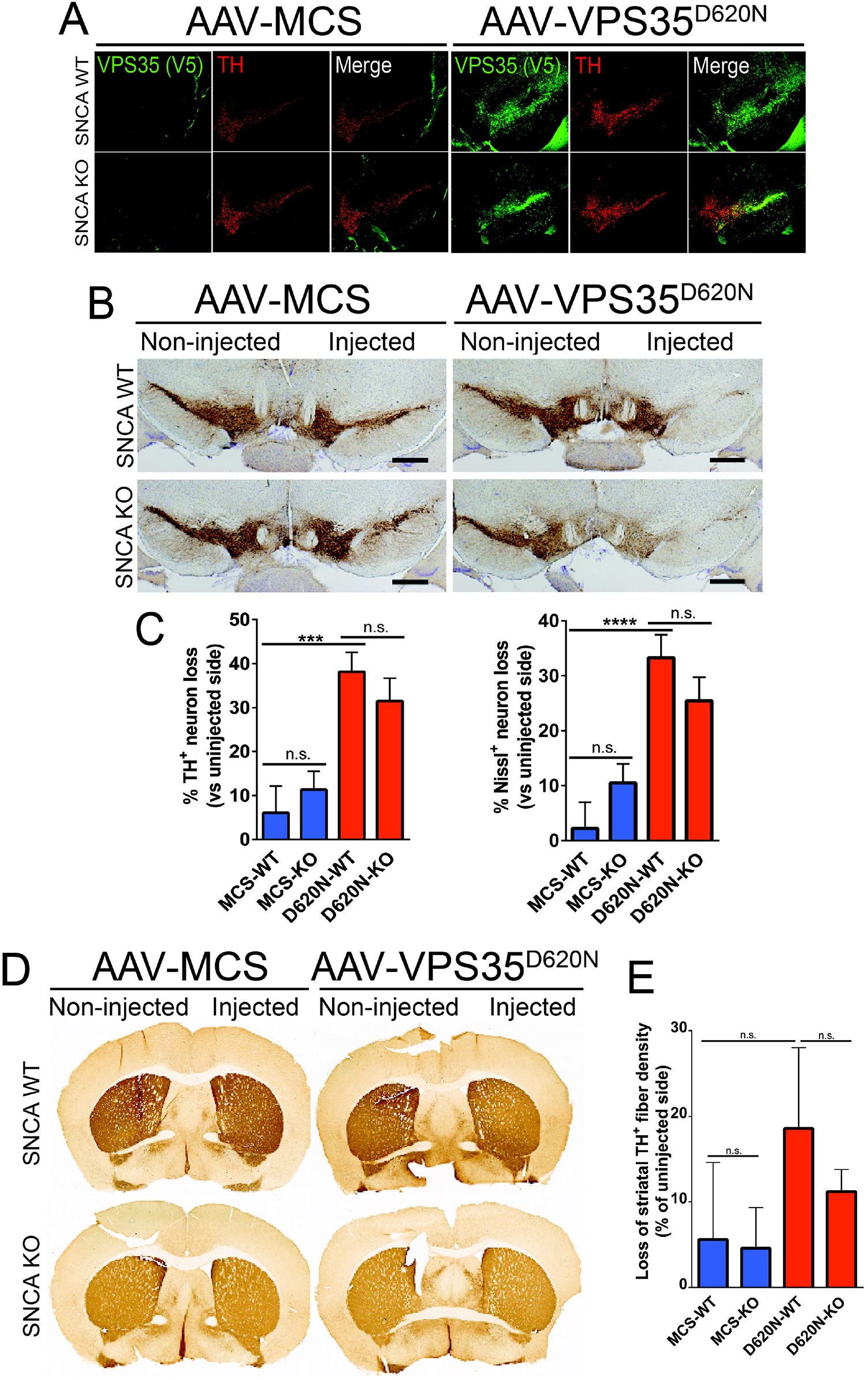
Dopaminergic neurodegeneration induced by human D620N VPS35 expression in mice occurs independent of endogenous _α_Syn. **A**) Immunofluorescent co-localization of human VPS35 (V5) with TH-positive neurons in the ipsilateral substantia nigra of WT or *SNCA* KO mice at 12 weeks following the injection of AAV2/6 vectors (control MCS or D620N VPS35). **B**) Immunohistochemical staining of nigral TH-positive neurons at 12 weeks after unilateral intranigral injection of MCS or D620N VPS35 vectors. **C**) Stereological quantitation of nigral TH-positive dopaminergic and total Nissl-positive neuronal loss in WT and *SNCA* KO mice induced by D620N VPS35 or control virus at 12 weeks. Data are expressed as percent neuronal loss relative to the uninjected nigra, with bars representing the mean ± SEM, *n* = 9-12 mice/group. \*\*\**P*<0.001 or \*\*\*\**P*<0.0001 by one-way ANOVA with Tukey’s multiple comparison test, as indicated. **D**) Representative photomicrographs of immunostaining for TH-positive nerve terminals in the striatum at 12 weeks following AAV vector delivery. **E**) Quantitation of striatal TH-positive fibers by optical density. Data are expressed as percent loss of TH-positive fibers relative to the uninjected side, with bars representing the mean ± SEM, *n* = 9-12 mice/group. n.s., non-significant by one-way ANOVA with Tukey’s multiple comparison test.

We next evaluated whether D620N VPS35 expression in the mouse nigra is sufficient to induce alterations in classic neuropathological markers using immunohistochemical analysis. D620N VPS35 expression (but not a control virus) induces the qualitative accumulation of phospho-Ser129-αSyn, a pathological form of αSyn, in the injected substantia nigra of WT mice relative to the contralateral nigra (**Fig. 3A**). D620N VPS35 fails to induce phospho-Ser129-αSyn immunostaining in *SNCA* KO mice, thereby confirming the specificity of this αSyn marker (**Fig. 3A**). Notably, immunoreactivity for total αSyn is not altered by D620N VPS35 expression in the nigra of WT mice (**Fig. 3A**), whereas *SNCA* KO brains lack signal for total αSyn as expected. We also evaluated immunostaining for amyloid precursor protein (APP), a sensitive marker of axonal damage (30, 35). Intriguingly, D620N VPS35 expression increases APP-positive staining in cell soma and neurites in the injected nigra of WT mice compared to the non-injected nigra, whereas this increase in APP staining is markedly attenuated in *SNCA* KO mice (**Fig. 3A**). This finding suggests that the increase in APP accumulation induced by D620N VPS35 is dependent on endogenous αSyn.

**Figure 3:**
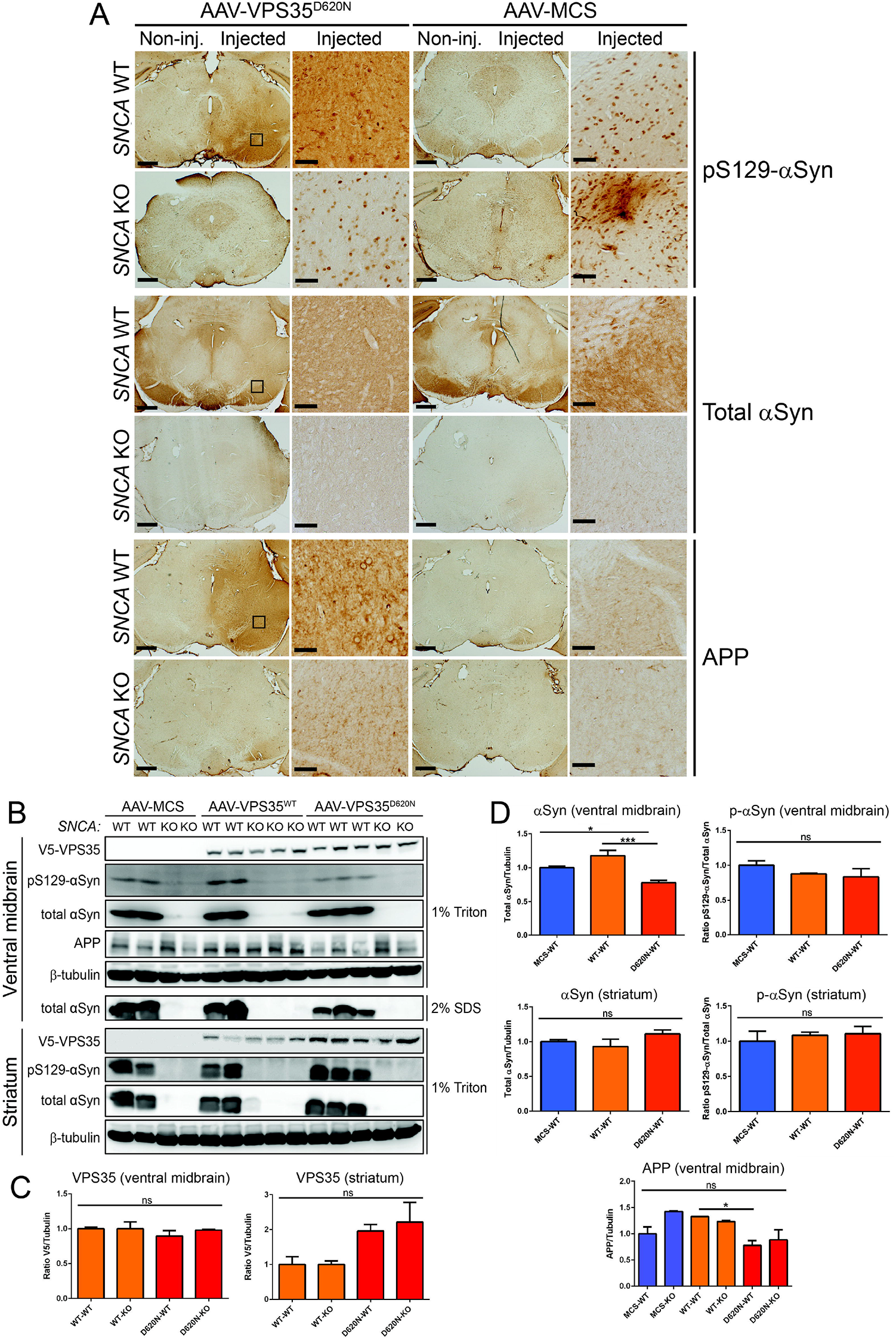
Accumulation of pSer129-_α_Syn and APP in the substantia nigra of WT mice induced by D620N VPS35 expression. **A**) Immunohistochemical staining of substantia nigra from WT and *SNCA* KO mice at 12 weeks after intranigral delivery of AAV2/6 vector expressing human D620N VPS35 (versus MCS) reveals the specific accumulation of pSer129-αSyn and APP in WT but not KO mice. Levels/distribution of total αSyn are normal in WT mice but absent from KO mice. Higher power images of the injected nigra (boxed) are shown. **B**) Western blot analysis of injected ventral midbrain and striatal extracts (1% Triton-X100 or 2% SDS fractions) from WT or *SNCA* KO mice at 12 weeks following the intranigral delivery of AAV2/6 vectors (MCS, WT VPS35 or D620N VPS35). Blots were probed with antibodies to V5 (VPS35), pS129-αSyn, total αSyn and APP, with β-tubulin as a loading control. **C-D**) Densitometric analysis of human VPS35 (V5), total αSyn, pS129-αSyn and APP levels in ventral midbrain and striatal extracts of WT or KO mice normalized to β-tubulin levels. pS129-αSyn levels were normalized to total αSyn. Bars represent mean ± SEM (*n* = 3-4 animals/group). \**P*<0.05 or \*\*\**P*<0.001 by one-way ANOVA with Dunnett’s *post-hoc* analysis, as indicated. n.s., non-significant.

To further evaluate alterations in these markers and to confirm equivalent levels of human VPS35 variants in these animals, Western blot analysis was conducted on Triton-soluble or SDS-soluble (Triton-insoluble) extracts from the injected ventral midbrain and striatum at 12 weeks post-injection. Here, we also delivered an AAV2/6 vector expressing human WT VPS35 for comparison. WT and D620N VPS35 are detected at equivalent levels in Triton-soluble ventral midbrain extracts of WT and *SNCA* KO mice (**Fig. 3B-C**). WT and D620N VPS35 are also detected in Triton-soluble striatal extracts, thereby confirming the efficient anterograde dopaminergic axonal transport of human VPS35 from the injected nigra to the striatum of WT and KO mice (**Fig. 3B-C**). Total αSyn and pSer129-αSyn levels are not increased by WT or D620N VPS35 expression compared to control virus in Triton-soluble ventral midbrain or striatal extracts from WT mice, and are absent from KO brains as expected (**Fig. 3B, D**). Instead, we observe a modest decrease in total αSyn levels in the ventral midbrain induced by D620N VPS35 relative to control virus or WT VPS35 (**Fig. 3B, D**). Full-length APP levels are also not increased in Triton-soluble ventral midbrain extracts expressing VPS35 variants, and instead APP levels are significantly reduced in WT mice by D620N VPS35 relative to the WT protein (**Fig. 3B, D**). While our biochemical data does not support an accumulation of pSer129-αSyn or full-length APP proteins induced by D620N VPS35 in WT mice (**Fig. 3B-D**), as suggested by immunohistochemical analyses (**Fig. 3A**), these data might suggest a redistribution of these pathological markers in neurons of the ventral midbrain in an αSyn-dependent manner. As such, D620N VPS35 expression in the mouse nigrostriatal pathway does not produce obvious αSyn neuropathology that would be consistent with the pathological aggregation of αSyn.

### Pathological _α_Syn does not cause a retromer deficiency in the mouse brain

Recent studies have suggested a role for the retromer in regulating the accumulation of αSyn in the hippocampus of transgenic mice expressing human WT αSyn, and the neuroprotective effects of viral-mediated VPS35 overexpression in this model (28). These studies suggest that pathological αSyn may mediate neurotoxicity by inducing a retromer deficiency. To explore this possibility, we first assessed retromer levels in prion promoter (PrP)-driven human A53T-αSyn transgenic mice that develop a progressive neurodegenerative phenotype characterized by neuronal degeneration in spinal cord and brainstem, reactive gliosis, αSyn aggregation, and motor deficits that lead to limb paralysis and premature lethality (between 7-15 months) (36). We evaluated the steady-state levels of the core retromer subunits, VPS35, VPS26 and VPS29, in four affected brain regions derived from 6 month-old asymptomatic (**Fig. 4A**) or end-stage symptomatic (**Fig. 4B-C**) A53T-αSyn mice and their non-transgenic littermates by Western blot analysis. We find that retromer levels are not significantly altered in the spinal cord, brainstem, ventral midbrain or striatum of 6 month-old and end-stage A53T-αSyn mice (**Fig. 4A-C**). Human αSyn expression and pSer129-αSyn are detectable in each brain region of the A53T-αSyn mice, as expected (**Fig. 4A-B**). Our data fail to provide evidence for the depletion of core retromer subunits in affected brain regions of A53T-αSyn mice with advancing age.

**Figure 4:**
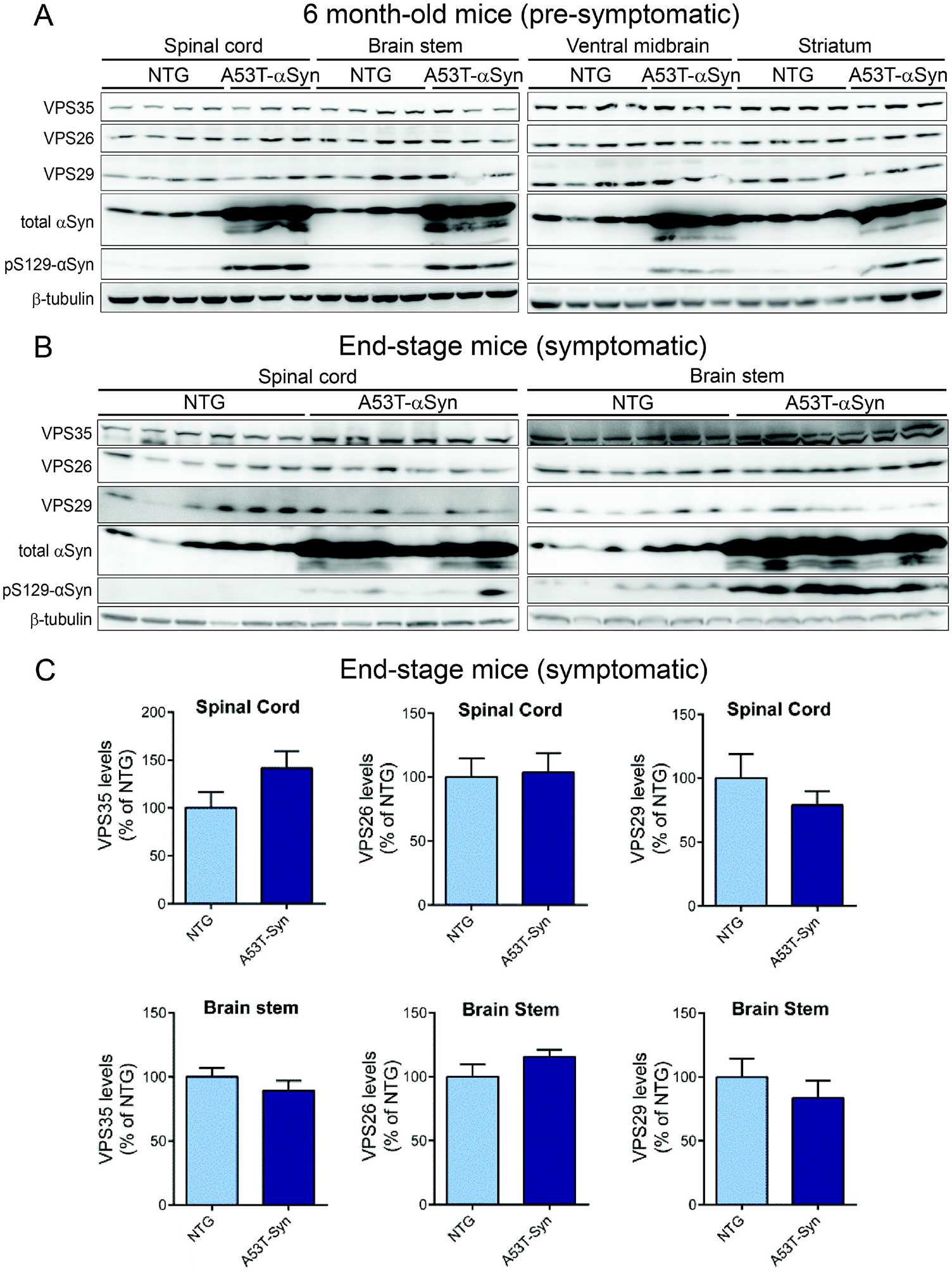
Absence of retromer deficiency in brains from human A53T-_α_-Syn transgenic mice. **A**) Western blot analysis of retromer subunits and αSyn (total or pS129) in Triton-soluble extracts from spinal cord, brain stem, ventral midbrain and striatum of 6-month old pre-symptomatic hemizygous A53T-αSyn transgenic mice and their non-transgenic littermates (*n* = 3-4 animals/group). **B**) Similar Western blot analysis of spinal cord and brain stem extracts from symptomatic end-stage hemizygous A53T-αSyn mice (∼12-13 months) and their non-transgenic littermates (*n* = 6 animals/group). **C**) Densitometric analysis of VPS35, VPS29 or VPS26 levels in the spinal cord and brain stem of end-stage A53T-αSyn transgenic and non-transgenic mice normalized to β-tubulin levels. Data are expressed as a percent of non-transgenic mice with bars representing the mean ± SEM (*n* = 6 animals/group). n.s., non-significant by unpaired two-tailed Student’s *t*-test.

### Retromer deficiency does not regulate _α_Syn pathology or premature lethality in A53T-_α_Syn transgenic mice

To further probe whether αSyn-induced neurotoxicity is mediated or exacerbated by retromer deficiency, we crossed heterozygous *VPS35* null mice with A53T-αSyn transgenic mice. We previously described viable heterozygous *VPS35*^*FLOX/WT*^ mice carrying a conditional allele (floxed “WT VPS35” mini-gene) that inadvertently disrupts normal *VPS35* expression and serves as a null allele (30). We find that heterozygosity for *VPS35* does not alter the steady-state levels of pSer129-αSyn or total αSyn in the striatum, ventral midbrain, brainstem or spinal cord of end-stage A53T-αSyn mice by Western blot analysis (**Fig. 5A-B**). The levels of VPS35 as well as VPS26 and VPS29 are markedly reduced in *VPS35*^*FLOX/WT*^ brains (**Fig. 5A**), as expected. We also examined the substantia nigra from 6 month-old pre-symptomatic mice for αSyn pathology using the pSer129-αSyn antibody, however, *VPS35* heterozygosity does not qualitatively alter the distribution, morphology or burden of pSer129-αSyn-positive pathology in A53T-αSyn mice (**Fig. 5C**). *VPS35*^*WT/WT*^ and *VPS35*^*FLOX/WT*^ mice lacking the A53T-αSyn transgene do not exhibit αSyn pathology (**Fig. 5C**), as expected. The impact of *VPS35* heterozygosity on the survival of A53T-αSyn mice was also monitored over 24 months. We find that A53T-αSyn mice display impaired survival and exhibit premature death owing to neurodegeneration over 10 to 18 months, with no significant effect of removing one *VPS35* allele (**Fig. 5D**). Finally, we conducted pilot studies to explore if *VPS35* heterozygosity influences the initial development of αSyn pathology induced by the unilateral intrastriatal delivery of mouse αSyn preformed fibrils (PFFs). At 30 days post-inoculation of PFFs, we detect pSer129-αSyn-positive pathology in the ipsilateral substantia nigra pars compacta that co-localizes with TH-positive dopaminergic neurons, yet the extent of pathology is qualitatively similar between *VPS35*^*WT/WT*^ and *VPS35*^*FLOX/WT*^ mice (**Fig. S1**). Our data indicate that retromer deficiency does not influence the lethal neurodegenerative phenotype of A53T-αSyn transgenic mice or the initial propagation of αSyn pathology in a PFF-based mouse model.

**Figure 5:**
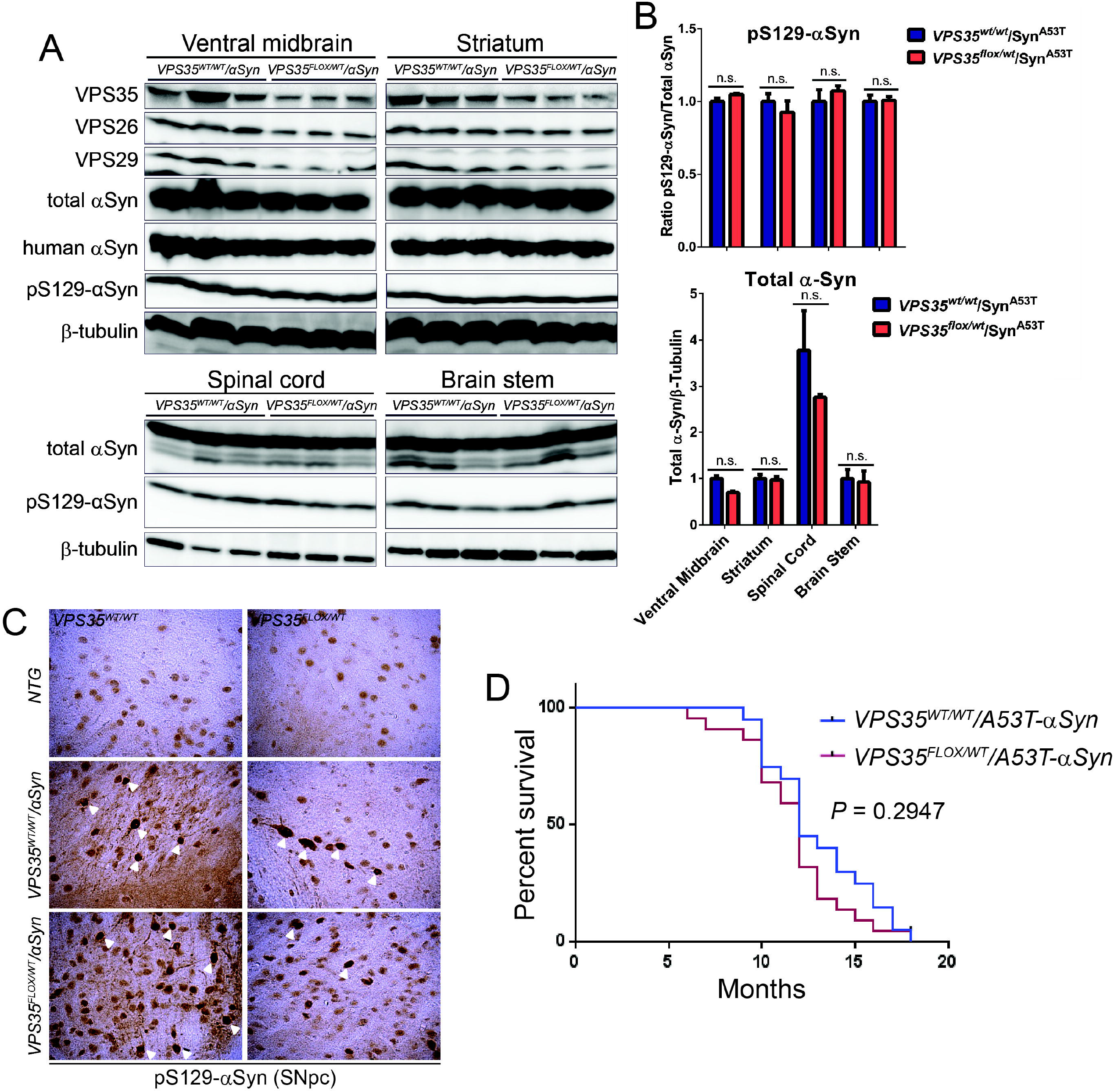
Heterozygous *VPS35* deletion fails to modify _α_-Syn levels and pathology or premature survival in human A53T-_α_-Syn transgenic mice. **A**) Western blot analyses of 1% Triton-soluble extracts from ventral midbrain, striatum, spinal cord or brain stem from ∼13-month old heterozygous *VPS35* null mice crossed with human A53T-α-Syn transgenic mice (*VPS35*^*FLOX/WT*^*/*_α_*Syn*) versus their A53T-α-Syn littermates (*VPS35*^*WT/WT*^*/*_α_*Syn*). Blots were probed for retromer subunits (VPS35, VPS26 and VPS29), total (Syn1), human (Syn211) and pathological (pS129-αSyn) α-synuclein or β-tubulin. **B**) Densitometric analysis of pS129-αSyn levels normalized to total αSyn, or total αSyn normalized to β-tubulin, expressed as a proportion of *VPS35*^*WT/WT*^/αSyn^A53T^ mice (mean ± SEM, *n* = 3 mice/genotype). *P*>0.05 by one-way ANOVA with Bonferroni’s *post-hoc* test, as indicated. n.s., non-significant. **C**) Representative images of immunohistochemical staining of pS129-αSyn in the substantia nigra (SNpc) of ∼6-month old pre-symptomatic *VPS35*^*WT/WT*^/αSyn^A53T^ or *VPS3*^*FLOX/WT*^/αSyn^A53T^ mice indicating equivalent α-Syn aggregation/accumulation (*white arrowheads*). Non-transgenic (NTG) littermate mice do not exhibit specific α-Syn pathology, as expected. **D**) Lethal neurodegenerative phenotype of human A53T-α-Syn transgenic mice is independent of VPS35 expression. Kaplan-Meier survival curves were generated by monitoring cohorts of *VPS35*^*WT/WT*^/αSyn^A53T^ (*n* = 27) and *VPS35*^*FLOX/WT*^/αSyn^A53T^ (*n* = 24) mice over 18 months until animals had to be euthanized due to the onset of terminal disease. There is no significant difference in survival between the two genotypes by log-rank (Mantel-Cox) test (*P* = 0.2947).

### Overexpression of wild-type VPS35 fails to protect against dopaminergic neuronal degeneration induced by human WT _α_Syn in the rat brain

We next sought to extend the recent intriguing observation by Dhungel *et al* demonstrating that the lentiviral-mediated overexpression of human WT VPS35 can attenuate hippocampal neuronal loss and αSyn pathology that develops in human WT-αSyn transgenic mice (28). We asked whether increasing VPS35 levels could similarly provide protection against dopaminergic neurodegeneration induced by human WT-αSyn. The AAV-mediated delivery of human WT-αSyn provides a more relevant rodent model of PD, compared to WT-αSyn transgenic mice, in that it recapitulates the progressive and robust degeneration of the nigrostriatal dopaminergic pathway and nigral αSyn pathology (37-39). Recombinant AAV2/6 vectors expressing human WT-αSyn and V5-tagged human VPS35 (WT or PD-linked D620N) or an empty control virus (MCS), were unilaterally co-injected into the substantia nigra of adult rats. At 14 weeks post-injection, we monitored dopaminergic neuronal degeneration, αSyn pathology and gliosis. Human VPS35 variants and human WT-αSyn could be robustly detected by immunohistochemistry in the ipsilateral rat substantia nigra and striatum for each animal group (**Fig. 6A-B**), and human VPS35 variants and αSyn substantially co-localize together in TH-positive nigral dopaminergic neurons by confocal immunofluorescence microscopy (**Fig. 6C**). To evaluate the impact of VPS35 overexpression on WT-αSyn-induced neurodegeneration, unbiased stereology was used to count TH-positive dopaminergic and total Nissl-positive neurons in the nigra (**Fig. 6D-E**). We find that WT-αSyn expression alone (αSyn + MCS) induces a ∼57% loss of dopaminergic neurons in the injected ipsilateral nigra of rats, and the co-expression of WT VPS35 has no impact on this robust neuronal loss (∼54%, **Fig. 6D-E**). Surprisingly, co-expression with D620N VPS35 markedly reduces αSyn-induced neuronal loss to ∼33%, but this effect is not significant relative to WT-αSyn alone (**Fig. 6D-E**). The expression of WT or D620N VPS35 alone at this lower viral titer induces ∼24 or 26% neuronal loss, respectively (**Fig. 6D-E**), comparable to our prior studies (32), whereas empty control virus produces negligible neuronal loss (**Fig. 6D**). We observe a comparable loss of total Nissl-positive nigral neurons by stereological counting (**Fig. 6E**), confirming dopaminergic neuronal degeneration. WT-αSyn expression also induces a marked ∼20% loss of TH-positive dopaminergic nerve terminals in the ipsilateral striatum, that parallels TH-positive neuronal loss, yet with no significant effect of co-expressing WT VPS35 and a non-significant reduction with D620N VPS35 (**Fig. 6F-G**). Together, these data indicate that WT VPS35 expression is not sufficient for neuroprotection against nigrostriatal dopaminergic neurodegeneration induced by WT-αSyn in this rat AAV model, whereas D620N VPS35 expression exhibits a modest protective effect.

**Figure 6:**
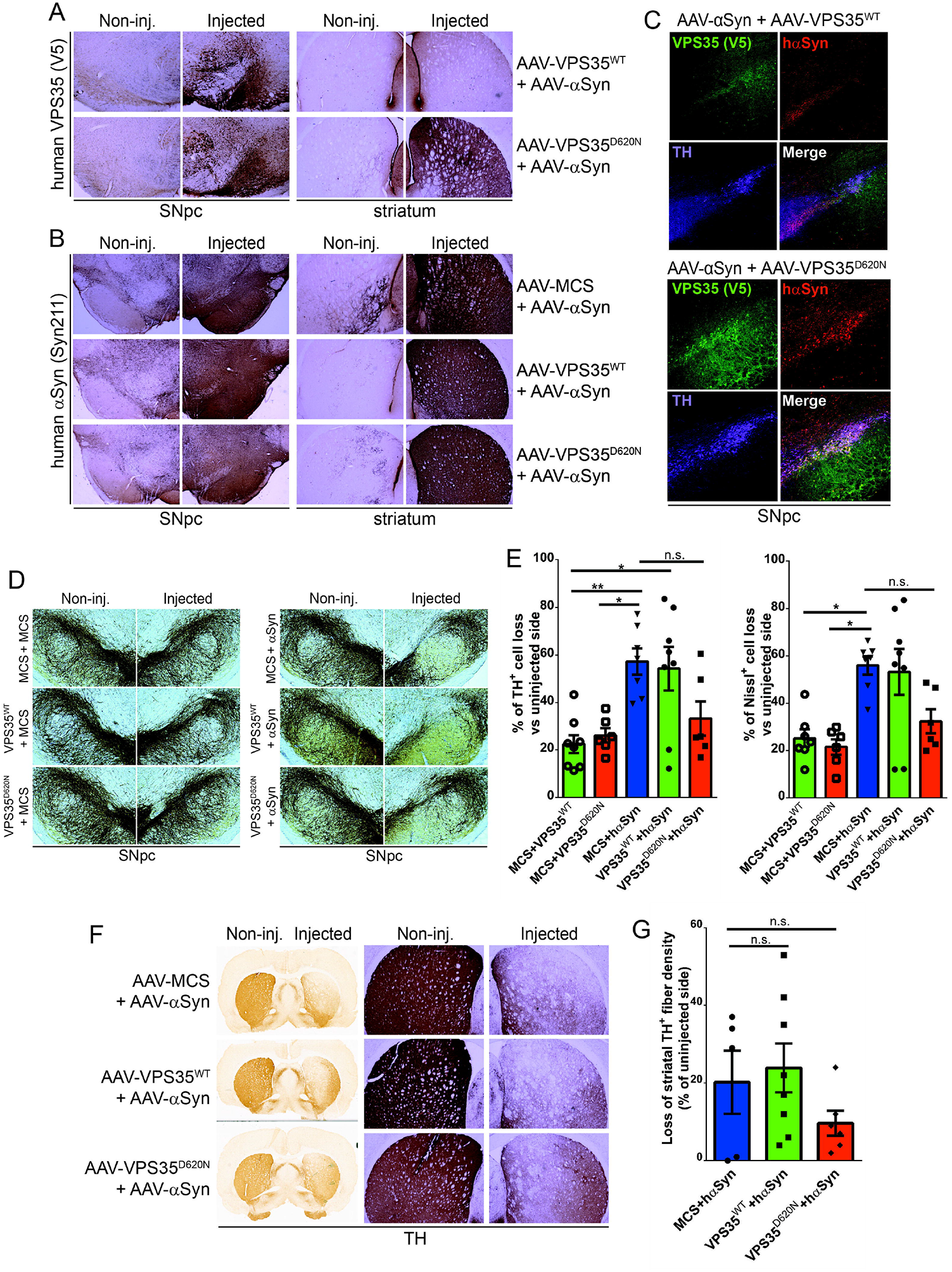
VPS35 overexpression fails to protect against nigrostriatal dopaminergic neurodegeneration induced by human WT-_α_-Syn in adult rats. **A-B**) Representative photomicrographs of (**A**) human VPS35 (V5) and (**B**) human α-synuclein (Syn211) immunostaining in the substantia nigra (SNpc) and striatum of adult rats at 14 weeks after co-injection of AAV2/6 vectors expressing WT-α-Syn together with VPS35 (WT or D620N) or empty control (MCS). Non-injected (left) and injected (right) hemispheres are indicated. **C**) Fluorescent immunostaining indicating marked co-localization of human VPS35 (WT or D620N) and human WT-α-Syn with TH-positive dopaminergic neurons of the injected rat substantia nigra at 14 weeks. **D**) Representative photomicrographs of anti-TH immunostaining in the rat substantia nigra (SNpc) at 14 weeks after the co-injection of AAV2/6 vectors expressing combinations of human VPS35 (WT or D620N), human WT-αSyn, or empty control (MCS). **E**) Unbiased stereological quantification of TH-positive dopaminergic and total Nissl-positive neurons in the substantia nigra at 14 weeks post-injection of AAV vectors. Data are expressed as percent cell loss relative to non-injected nigra with bars indicating the mean ± SEM (*n* = 7-8 animals/group). \**P*<0.05 or \*\**P*<0.01 by one-way ANOVA with Tukey’s multiple comparison test, as indicated. n.s., non-significant. **F**) Representative photomicrographs of immunostaining for TH-positive nerve terminals in the striatum at 14 weeks after AAV vector delivery in injected and non-injected hemispheres. **G**) Quantitation of optical density of TH-positive immunostaining in the striatum. Data are expressed as percent loss of TH-positive fibers relative to the non-injected side, with bars indicating the mean ± SEM (*n* = 7-8 animals/group). n.s., non-significant by one-way ANOVA with Tukey’s multiple comparison test.

We next sought to evaluate the impact of human VPS35 expression on αSyn pathology and gliosis in this rat model. Extracts derived from the ventral midbrain and striatum of AAV-αSyn/VPS35 co-injected rats at 14 weeks post-injection were analyzed by Western blot. Human WT and D620N VPS35 proteins are detected in Triton-soluble fractions of ventral midbrain and striatum, with a non-significant increase of WT VPS35 compared to D620N protein (**Fig. 7A-C**). The steady-state levels of Triton-soluble or SDS-soluble human αSyn are not altered by co-expression with D620N VPS35 relative to αSyn alone (+ MCS) in both brain regions, whereas co-expression of WT VPS35 leads to a significant increase of Triton-soluble human αSyn in the striatum and a non-significant increase in ventral midbrain (**Fig. 7A-C**). However, the levels of pSer129-αSyn are significantly reduced by co-expression with WT VPS35 in the Triton-soluble striatum with a non-significant reduction in ventral midbrain, whereas D620N VPS35 co-expression also leads to a modest non-significant reduction of pSer129-αSyn in both brain regions (**Fig. 7A-C**). We further evaluated the impact of VPS35 on αSyn pathology and αSyn-induced reactive gliosis using immunohistochemistry in the rat substantia nigra at 14 weeks post-injection. Phospho-Ser129-αSyn-positive pathology is robustly detected localized to neuronal soma and neurites in the ipsilateral substantia nigra pars compacta of rats injected with AAV-αSyn alone but is absent from the non-injected nigra (**Fig. 8A-B, Q**). The co-expression of D620N VPS35 induces a significant increase in pSer129-αSyn pathology with quantitation using Halo analysis software, compared to AAV-αSyn alone, whereas WT VPS35 produces an intermediate effect (**Fig. 8C-D, Q** and **Fig. S2**). The expression of human αSyn alone fails to induce a significant increase in the area occupied by GFAP-positive astrocytes or Iba1-positive microglia, relative to the non-injected contralateral nigra, and the co-expression with WT or D620N VPS35 has no effect (**Fig. 8E-L, R, S** and **Fig. S2**). We also monitored activated microglia based on the morphology of Iba1-positive microglia using Halo software (**Fig. 8T** and **Fig. S2**) or CD68-positive microglia (**Fig. 8M-P, U** and **Fig. S2**), yet human αSyn expression alone or its co-expression with VPS35 variants induces only modest yet non-significant microglial activation at 14 weeks in this AAV model. Collectively, these data fail to convincingly demonstrate a neuroprotective effect of WT VPS35 overexpression in attenuating human αSyn pathology or gliosis. Instead, we observe a biochemical increase in human αSyn levels induced by WT VPS35, or an increase in pSer129-αSyn-positive pathology induced by D620N VPS35 despite its protective effects on dopaminergic neurodegeneration.

**Figure 7:**
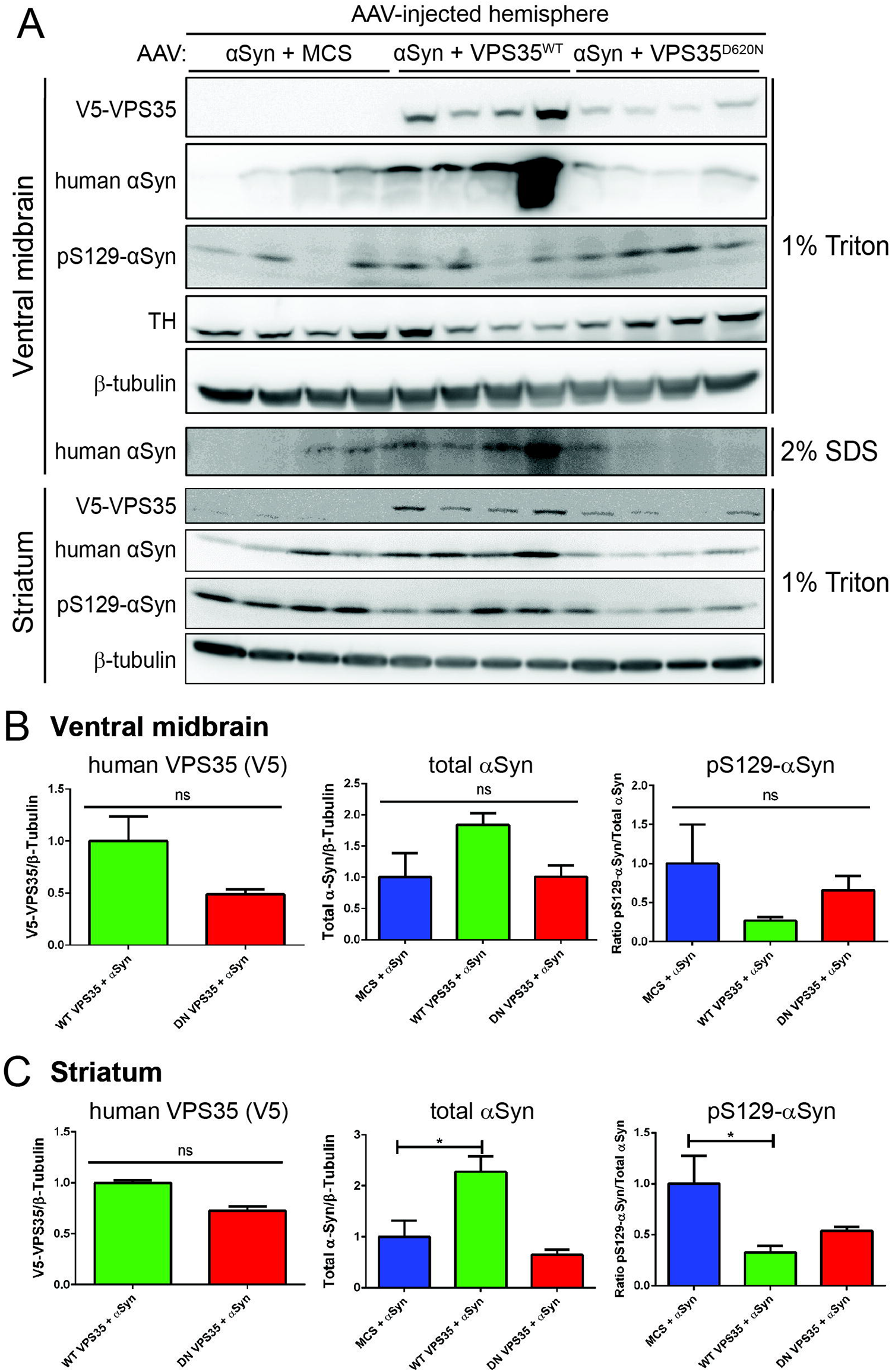
Biochemical effect of VPS35 overexpression on human WT-_α_-Syn levels and phosphorylation in rat brain. **A**) Western blot analysis of rat ventral midbrain and striatal extracts at 14 weeks post-injection of AAV vectors expressing WT-α-Syn and VPS35 (WT or D620N) or empty control (MCS). 1% Triton-X100 or 2% SDS fractions were probed with antibodies to human VPS35 (V5), human α-Syn (Syn211 antibody), pS129-α-Syn, TH, or β-tubulin as a control. Note, only human WT-α-Syn is detected in the 2% SDS fraction from ventral midbrain, but pS129-α-Syn or human VPS35 are not detectable. **B-C**) Densitometric analysis of pS129-α-Syn levels normalized to human α-Syn, or human α-Syn or VPS35 (V5) levels normalized to β-tubulin in 1% Triton-soluble extracts from (**B**) ventral midbrain or (**C**) striatum. Bars represent mean ± SEM (*n* = 4 animals/group). \**P*<0.05 by one-way ANOVA with Dunnett’s *post-hoc* analysis, as indicated. n.s., non-significant.

**Figure 8:**
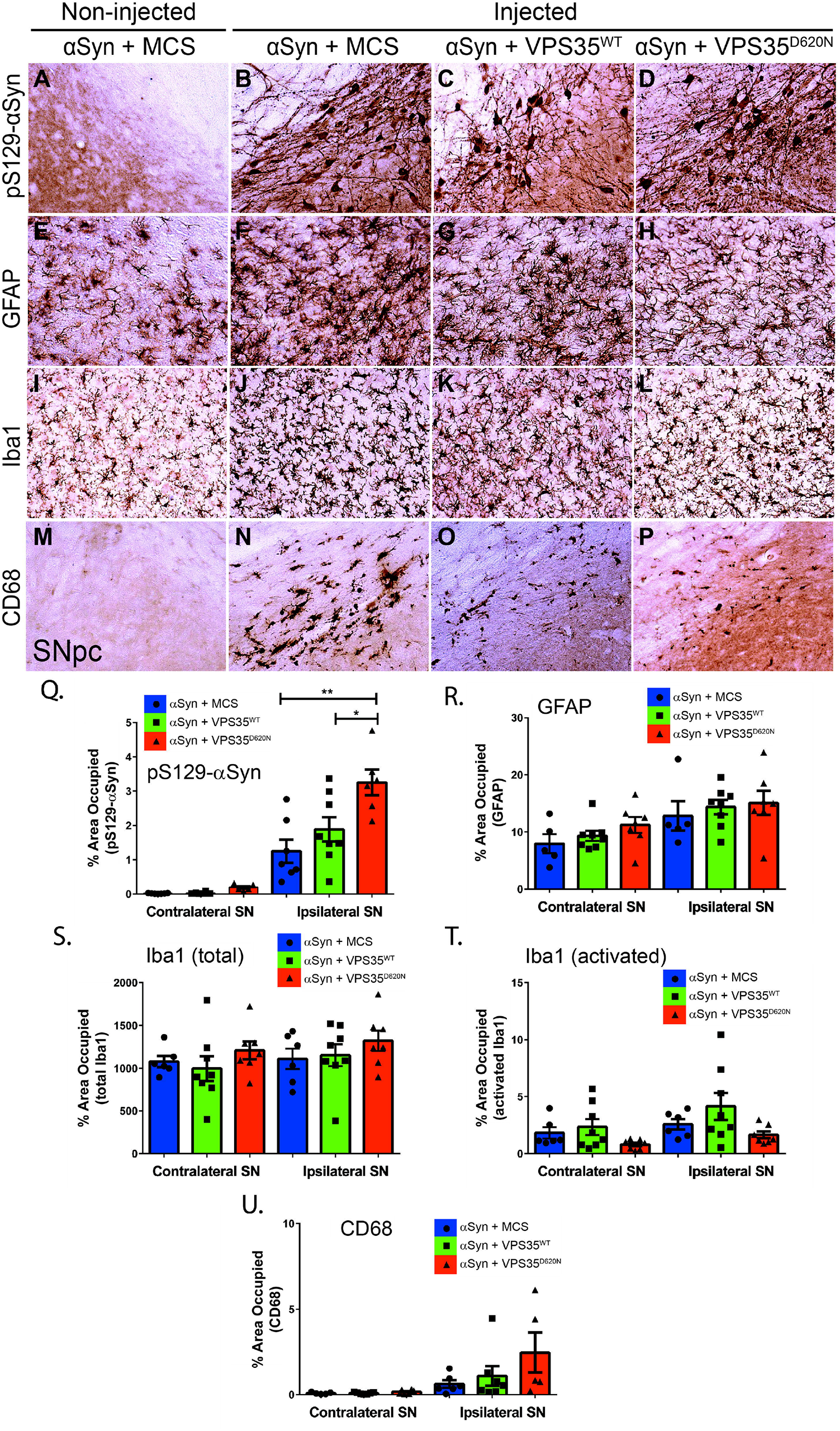
VPS35 overexpression does not attenuate human WT-_α_-Syn-dependent pathology in the rat substantia nigra. **A-P**) Immunohistochemical staining of rat substantia nigra pars compacta (SNpc) at 14 weeks following intranigral co-injection of AAV vectors expressing human WT-α-Syn with human VPS35 (WT or D620N) or empty control (MCS). Immunoreactivity for (**A-D**) pS129-α-Syn, (**E-H**) astrocytes (GFAP-positive), (**I-L**) total microglia (Iba1-positive), and (**M-P**) activated microglia (CD68-positive), in the injected SNpc are shown. The non-injected SNpc from the α-Syn/MCS control group is shown for comparison. **Q-U)** Quantitation of each pathological marker using HALO analysis software. SNpc sections with pS129-α-Syn, GFAP, Iba1 (total or activated) and CD68 immunostaining were quantified as the percent area occupied by each marker in the ipsilateral (injected) or contralateral (non-injected) hemispheres. Activated Iba1-positive microglia were identified from total microglia based on morphological criteria using Halo (**T**). Bars represent mean ± SEM (*n* = 5-8 animals/group). \**P*<0.05 or \*\**P*<0.01 by one-way ANOVA with Tukey’s multiple comparison’s test, as indicated.

## Discussion

Here, we have investigated the biochemical, functional and pathological interaction of VPS35 and αSyn in cells and rodent brain. We fail to identify a robust biochemical interaction between VPS35 and αSyn, and limited effects of αSyn variant overexpression on retromer levels, whereas VPS35 depletion fails to induce the accumulation of αSyn in human cells. In the mouse brain, we find that the robust degeneration of the nigrostriatal dopaminergic pathway induced by viral-mediated human D620N VPS35 expression is not robustly altered by the deletion of endogenous αSyn. D620N VPS35 expression does lead to an increase of pSer129-αSyn and full-length APP immunoreactivity in the substantia nigra yet this effect is not confirmed in brain extracts, potentially implying a redistribution of these markers rather than their accumulation. In a well-characterized human A53T-αSyn transgenic mouse model that develops a lethal neurodegenerative phenotype, we find no evidence for a physical retromer deficiency in multiple brain regions affected by αSyn pathology. Furthermore, *VPS35* heterozygosity has no impact on αSyn neuropathology or the lethal neurodegenerative phenotype that occurs in this A53T-αSyn model. Finally, we find that the viral-mediated overexpression of human WT VPS35 does not protect against nigrostriatal dopaminergic pathway degeneration, αSyn pathology or reactive gliosis induced by the overexpression of human WT αSyn in the rat substantia nigra. Our data suggest that endogenous αSyn expression is not critically required for dopaminergic neurodegeneration induced by human D620N VPS35, and furthermore, we find no evidence that mutant αSyn induces a retromer deficiency or that restoring VPS35 is sufficient to mediate neuroprotection against αSyn-induced neurotoxicity. Taken together, our study fails to provide compelling support for a functional or pathological bidirectional interaction between VPS35 and αSyn in mediating neurodegeneration in the rodent brain, suggesting that both proteins may operate via distinct pathways.

Recent studies in mouse models with a germline heterozygous *VPS35* deletion (22), or conditional homozygous *VPS35* deletion selectively in nigral dopaminergic neurons (25), suggest that reducing VPS35 levels causes an accumulation of total αSyn in neurons. However, αSyn pathology as defined by the presence of pSer129-αSyn or αSyn aggregation has not been reported in the brain of these knockout models (22, 25). Similarly, *D620N VPS35* knockin mice do not exhibit endogenous αSyn pathology (30), or the capacity to modulate the lethal neurodegenerative phenotype of human A53T-αSyn transgenic mice (30), but there is some evidence for the modest accumulation of total αSyn in the brain (31). Despite the absence of obvious αSyn aggregation, it has been suggested that neuronal αSyn accumulation in these knockout or knockin models could contribute to neurodegeneration (22, 25, 31). We have previously reported that the AAV-mediated expression of human D620N VPS35 in the substantia nigra of adult rats is sufficient to induce the degeneration of dopaminergic neurons yet this occurs in the absence of obvious αSyn pathology (32). Here, we extend these studies to mice to address a functional requirement of endogenous αSyn for mediating the pathogenic effects of D620N VPS35. While we find evidence that D620N VPS35 can specifically induce pSer129-αSyn immunoreactivity in the ipsilateral substantia nigra of WT mice (**Fig. 3A**), that is absent from *SNCA* KO mice as expected and thus confirms the specificity of this signal, we do not observe a robust impact of αSyn removal on dopaminergic neuronal loss in this AAV model (**Fig. 2**). Our study implies that while VPS35 can regulate pSer129-αSyn levels in neuronal soma and processes of the ventral midbrain, this does not meaningfully contribute to the degenerative effects of D620N VPS35, and therefore it is likely benign and does not represent a critical downstream pathogenic event. Additional studies are now needed in *VPS35* knockout or *D620N VPS35* knockin mice crossed to *SNCA* KO mice, to determine whether endogenous αSyn can contribute to neurodegenerative phenotypes that manifest in these models.

It is unclear how *VPS35* deletion or PD-linked mutations could drive the accumulation of αSyn in these mouse models, since αSyn is not a known retromer cargo and our studies in human cells do not support the physical interaction of both proteins (**Fig. 1**). It is most likely that an effect of VPS35 on αSyn accumulation is indirect and may result from an impairment of lysosomal degradation due to the abnormal retromer sorting of CI-M6PR that delivers lysosomal enzymes such as cathepsin D to the lysosomal lumen (20, 21, 40), or via impaired sorting of the CMA receptor, LAMP2A (22). While modulating VPS35 can modestly regulate the levels of αSyn in the brain (22, 25, 31, 40), yet does not manifest classical αSyn pathology, VPS35 has been convincingly shown to regulate the aggregation-prone microtubule-associated protein tau (19). Several studies indicate that the levels of retromer subunit proteins are reduced in affected brain regions of human tauopathies (Progressive Supranuclear Palsy and Pick’s disease) and Alzheimer’s disease (41, 42), and silencing VPS35 expression in the brain can exacerbate tau neuropathology, and motor and learning impairments in human P301S-tau transgenic mice (42). Oppositely, retromer stabilization using the pharmacological chaperone TPT-172 (R33) can reduce Aβ deposition and abnormal tau, and improve memory impairments in the 3xTg mouse model of AD (43). In PD-linked *D620N VPS35* knockin mice that reveal a lack of αSyn pathology, these mice exhibit instead a widespread accumulation of abnormal somatodendritic tau in the brain characterized by pathological hyperphosphorylation and conformational-specific epitopes (30). Therefore, while VPS35 appears to have the capacity to regulate a number of protein aggregation pathways (19), our observations suggest that the accumulation of αSyn may not play a critical role in rodent models of PD. It will be important in future studies to similarly address the contribution of tau pathology to neurodegeneration induced by PD-linked *VPS35* mutations in such rodent models.

A number of studies have shown that reducing *VPS35* expression can exacerbate the toxicity associated with human αSyn in yeast, *C*.*elegans* and *Drosophila* models of PD (27, 28). It is unclear whether the effects of reducing *VPS35* are specific to αSyn-dependent pathways or act more generally in lowering cellular health and viability. Additional studies suggest that increasing VPS35 expression in mouse hippocampal neurons is sufficient to mediate neuroprotection in transgenic mice expressing human WT αSyn (line 61) (28). These studies combined tend to support the concept that human αSyn induces a retromer deficiency, with *VPS35* deletion exacerbating the pathogenic effects of αSyn and increased VPS35 expression being protective. How this mechanism is related to vulnerable neurons in PD is less clear. In our attempt to recreate and extend prior studies, we find that human A53T-αSyn transgenic mice do not exhibit reduced levels of retromer subunits in affected midbrain and hindbrain regions from mice that were pre-symptomatic or with end-stage disease despite a high burden of αSyn pathology (**Fig. 4**). Consistent with this finding, the germline heterozygous deletion of *VPS35* did not exacerbate αSyn accumulation, pathology and impaired survival that is characteristic of this A53T-αSyn mouse model (**Fig. 5**). One key difference with prior studies is that homozygous *VPS35* deletion in these lower organisms is not lethal as in mice, and our studies were therefore restricted to using heterozygous *VPS35* null mice that may not have been sufficient to exacerbate the pathogenic effects of A53T-αSyn. While our data does not support the findings reported in lower organisms suggesting that the retromer may operate downstream of αSyn, it is possible that the most relevant neuronal populations or brain regions are not adequately captured in our mouse studies. Human A53T-αSyn transgenic mice develop limb paralysis and impaired survival due mainly to the degeneration of spinal cord motor neurons yet these mice do exhibit widespread neuronal αSyn pathology (36, 44), however, we did not evaluate retromer levels in forebrain structures or specifically in individual neuronal populations such as nigral dopaminergic neurons. Nevertheless, we find no evidence that human A53T-αSyn drives a physical or functional retromer deficiency in this particular transgenic mouse model. It will be of interest to determine whether transgenic models based upon human WT αSyn (i.e. line 61) or driven by endogenous (i.e. BAC mice) rather than ectopic promoters display alterations in retromer subunit levels in relevant brain regions.

We further extended prior studies, reporting the protective effects of VPS35 overexpression against WT αSyn in the hippocampus of transgenic mice (28), to a well-characterized and robust rat model of PD based upon the AAV-mediated delivery of human WT αSyn (37, 38). This allowed us to evaluate the neuroprotective effects of VPS35 directly in the nigrostriatal dopaminergic pathway. We find that the overexpression of WT VPS35 has minimal impact on nigral dopaminergic neuronal loss, αSyn pathology and reactive gliosis induced by human WT αSyn expression (**Fig. 6-8**). The lack of protective effect of WT VPS35 in our αSyn rat model could reflect differences from prior studies between species (rat versus mouse), the susceptibility of distinct neuronal populations (nigral dopaminergic versus hippocampal pyramidal neurons), or inherent technical or mechanistic differences between transgenic and viral-mediated human αSyn expression. For example, VPS35 is prominently detected in rodent hippocampal neurons where it has been linked to the accumulation of tau pathology (30, 41, 42, 45). Notably, we do find a modest protective effect of D620N VPS35 against αSyn-induced neurotoxicity (**Fig. 6D-G**), supporting the concept that this PD-linked mutation might act through a gain-of-function mechanism due to increased or altered activity (19, 46). While we do observe increased nigral αSyn pathology induced by D620N VPS35 in this model (**Fig. 8**), this effect most likely correlates with the lower level of dopaminergic neuronal loss in this group (**Fig. 6E**). Our studies demonstrate that the degeneration of a PD-relevant neuronal population induced by human WT αSyn most likely is not due to a deficiency in VPS35 expression. It is possible that dopaminergic neurons do have reduced retromer activity following αSyn expression, but which requires the restoration of multiple subunits of the cargo-selective complex. Future studies might address a role for the retromer in αSyn-induced neurotoxicity by administering pharmacological chaperones such as R55 or R33 to these rodent models (47), in order to stabilize and increase the entire retromer complex rather than individual subunits. Our studies serve an important role in establishing that restoring WT VPS35 levels alone is not sufficient to protect dopaminergic neurons from human αSyn-induced neurotoxicity, in contrast to the effects reported in hippocampal pyramidal neurons (28).

Collectively, our data fail to provide robust support for a bidirectional pathway between VPS35 and αSyn in the brain using multiple well-characterized rodent models of PD. We demonstrate that endogenous αSyn is not critically required for the neurodegenerative effects of human D620N VPS35 in mice, and that modulating VPS35 levels via deletion or overexpression has minimal effects on two distinct and robust rodent models of human αSyn-induced neurodegeneration (A53T-αSyn transgenic mice or rats injected with AAV-WT-αSyn). To provide further clarity, future studies are now required in *post-mortem* human brains to evaluate whether Lewy pathology is a major feature of *VPS35*-linked PD (7), and furthermore, whether *SNCA*-linked PD brains exhibit reduced retromer subunits in affected regions or neurons similar to observations in AD, tauopathy and ALS brains (19, 33, 41, 42).

## Materials and Methods

### Animals

All animal experiments were approved by the Van Andel Institute Institutional Animal Care and Use Committee (IACUC) and conducted in strict accordance with the NIH Guild for the Care and Use of Laboratory Animals. Rodents were provided with food and water *ad libitum*, exposed to a 12 h light/dark cycle and maintained in a pathogen-free barrier facility. Female adult Sprague-Dawley rats (weighing ∼180-200 grams) were obtained from Charles River Laboratories and used for the stereotaxic delivery of AAV vectors. *VPS35*^*FLOX/WT*^ mice carrying a floxed “WT mini-gene” insertion that disrupts VPS35 expression (*Vps35*^*tm1*.*2Mjff*^, stock no. 021807) were obtained from The Jackson Laboratory and described previously (30). *SNCA* knockout mice (with deletion of exons 1-2; *Snca*^*tm1Rosl*^, stock no. 003692) were obtained from The Jackson Laboratory, and human A53T-α-Syn transgenic mice (line G2-3, driven by a mouse prion protein promoter; stock no. 006823) have been described (36). Mice were identified by genomic PCR using established genotyping protocols. Double mutant mice (A53T-α-Syn^Tg/+^/*VPS35*^*FLOX/WT*^) and appropriate littermate controls were generated by a single round of crossbreeding between hemizygous human A53T-α-Syn and heterozygous *VPS35*^*FLOX/WT*^ mice.

### Plasmids and Antibodies

A Myc-tagged human WT α-synuclein plasmid was described previously (48), and untagged human α-synuclein (WT, A30P, E46K, A53T) plasmids were obtained from Prof. Hilal Lashuel (EPFL, Switzerland). A Myc-tagged human tau (4R0N isoform, WT) plasmid was provided by Prof. Leonard Petrucelli (Mayo Clinic Jacksonville, Florida). GIPZ lentiviral plasmids co-expressing turboGFP and miR30-adpated short hairpin RNAs targeting VPS35 (shRNA #1, clone Id: V3LHS_389029; shRNA #2, clone Id: V2LMM_36638) or a non-silencing control (Dharmacon Cat # RHS4346) from a Pol II human CMV promoter were obtained from Horizon Discovery.

The following primary antibodies were used: mouse anti-V5, anti-V5-FITC and anti-V5-HRP (Life Technologies), mouse anti-Myc-HRP (clone 9E10; Roche), mouse anti-VPS35 (clone 2D3, ab57632; Abcam), rabbit anti-VPS26 (ab23892; Abcam), goat anti-VPS29 (ab10160; Abcam), rabbit anti-TH (NB300-109; Novus Biologicals), mouse anti-α-synuclein (clone 42; BD Biosciences), mouse anti-human α-synuclein (clone Syn211; Sigma), mouse anti-pSer129-α-synuclein (clone EP1536Y; Abcam), mouse anti-APP (clone 22C11; Millipore), rabbit anti-GFAP (ab227761; Abcam), rabbit anti-Iba1 (019-19741; Wako), mouse anti-rat CD68 (clone ED1; Bio-Rad), and mouse anti-β-tubulin (clone TUB 2.1; Sigma). Secondary HRP-conjugated antibodies used for Western blotting were: goat anti-mouse IgG, light chain-specific and mouse anti-rabbit IgG, light chain-specific (Jackson Immunoresearch). For fluorescence confocal analysis, the following secondary antibodies were used: AlexaFluor-488, -594 or -647 goat anti-mouse IgG, and AlexaFluor-488 or -647 goat anti-rabbit IgG (ThermoFisher). For bright-field microscopy, the following biotinylated secondary antibodies were used: goat anti-mouse IgG and goat anti-rabbit IgG (Vector Labs).

### Cell culture and transient transfection

Human SH-SY5Y neural cells and HEK-293T cells were maintained at 37°C with 5% CO_2_ in Dulbecco’s modified Eagle’s medium (DMEM) (Life Technologies) supplemented with 10% fetal bovine serum and penicillin/streptomycin. Transient transfection was achieved by transfecting cells with plasmid DNAs using XtremeGene HP DNA Transfection reagent (Roche) according to the manufacturer’s instructions. Cells were harvested 48-72 h post-transfection. To create cell lines stably expressing shRNAs, SH-SY5Y cells were transiently transfected with pGIPZ-turboGFP/shRNA expression plasmids as above followed by selection at 48 h post-transfection in media containing puromycin (2 µg/ml) for 2-3 weeks to produce stable clones.

### Co-immunoprecipitation assays and Western blotting

For co-immunoprecipitation (Co-IP) assays, HEK-239T cells were transiently co-transfected with plasmid combinations in 10 cm dishes. At 48 h post-transfection, cells were harvested in 1 ml of lysis buffer (20□mM HEPES-KOH, pH 7.2, 50□mM potassium acetate, 200□mM sorbitol, 2_mM EDTA, 0.1% Triton X-100, 1X Complete Mini protease inhibitor cocktail [Roche Applied Sciences]), allowed to rotate for 1 h at 4°C, and centrifuged at 15,000 rpm for 15 min. Soluble fractions were combined with Protein G-Dynabeads (Thermo Fisher) that had been pre-incubated with mouse anti-V5 antibody (2 µg; Life Technologies) and incubated overnight at 4°C. Dynabead complexes were washed once with 1X PBS, 0.1% Triton X-100, 150 mM NaCl, twice with 1X PBS, 0.1% Triton X-100, and three times with 1X PBS. IPs were eluted in Laemmli sample buffer by boiling at 95ºC for 5 min. IPs and input lysates (1% total) were resolved by SDS–PAGE, transferred to nitrocellulose (0.2 µm; GE Healthcare), and subjected to Western blotting with primary antibodies and light chain-specific anti-mouse/rabbit IgG-HRP conjugate antibodies. Proteins were visualized by enhanced chemiluminescence (ECL, GE Healthcare) and digital images were acquired using a FujiFilm LAS-4000 Image Analysis system. Images were subjected to densitometric quantitation using Image Studio Lite (LI-COR Biosciences).

### Recombinant AAV2/6 virus production

V5-tagged human VPS35 (WT or D620N) cDNAs or a stuffer sequence (WPRE) inserted into a pAAV2-mPGK-MCS vector were described previously (32). Recombinant AAV2/6 viral vectors were produced and titered as previously described (34) by the University of North Carolina Viral Vector Core. Viruses were diluted to a final concentration of ∼1.3 × 10^12^ viral genomes (vg) per ml. AAV2/6-PGK-αSyn-WT-WPRE or AAV2/6-PGK-MCS-WPRE vectors were provided by Dr. Bernard Schneider (Bertarelli Foundation Gene Therapy Platform, EPFL, Switzerland) and described previously (38), and were diluted to a viral titer of ∼1 × 10^10^ transducing units (TUs) per ml.

### Stereotactic surgeries

Stereotactic injections were performed as previously described (32, 34, 38). Briefly, *SNCA* KO or WT littermate mice aged 3-4 months received unilateral injections of AAV2/6-VPS35 or AAV2/6-MCS-WPRE vectors into the substantia nigra using the following coordinates relative to the bregma: anterior-posterior (A-P), -2.9 mm; medio-lateral (M-L), -1.3 mm; dorso-ventral (D-V), -4.2 mm. Each mouse received a viral titer of ∼2.6 × 10^9^ vg of AAV2/6 in a volume of 2 μl at a flow rate of 0.2 μl/minute. Animals were sacrificed at 12 weeks post-injection.

Female adult Sprague-Dawley rats (180-200 g) received a single unilateral intranigral co-injection of the following recombinant AAV2/6 vectors with transgene expression under the control of a strong ubiquitous PGK1 promoter: 1) AAV-PGK-MCS-WPRE x2, 2) AAV-PGK-αSyn-WT-WPRE + AAV-PGK-MCS-WPRE, 3) AAV-PGK-VPS35-WT + AAV-PGK-MCS-WPRE, 4) AAV-PGK-VPS35-D620N + AAV-PGK-MCS-WPRE, 5) AAV-PGK-αSyn-WT-WPRE + AAV-PGK-VPS35-WT, 6) AAV-PGK-αSyn-WT-WPRE + AAV-PGK-VPS35-D620N. AAV-αSyn-WT-WPRE or AAV-MCS-WPRE vectors were delivered at a titer of ∼1 × 10^7^ transducing units (TUs) in combination with AAV-VPS35 or AAV-MCS-WPRE vectors at a titer of ∼2 × 10^9^ vg. A total volume of 4 μl virus was injected unilaterally in the rat substantia nigra at the following coordinates: A-P, -5.2 mm; M-L, -2.0 mm; and D-V, −7.8 mm (relative to bregma). Animals were sacrificed at 14 weeks post-injection.

### Biochemical analysis of tissues

Rodent brain tissues were rapidly microdissected and homogenized in 6x volumes of Triton lysis buffer (50□mM Tris-HCl, pH 7.5, 150 mM NaCl, 5% glycerol, 1% Triton X-100, 1□mM EDTA, 1X Complete Mini protease inhibitor cocktail [Roche]). Tissues were disrupted using a mechanical homogenizer (IKA T10 basic, Ultra Turrax), and the Triton-soluble fraction was obtained after ultracentrifugation at 100,000 x *g* for 30 min at 4°C. Pellets were further extracted by sonication in 3x volumes of SDS lysis buffer (50□mM Tris-HCl, pH 7.4, 2% SDS, 1X Complete Mini protease inhibitor cocktail [Roche]) with centrifugation at 21,000 x *g* for 30 min at 25°C to obtain the SDS-soluble fraction (Triton-insoluble). Protein concentration was determined using the BCA assay (Pierce Biotech).

### Immunohistochemistry and Immunofluorescence

Rodents were transcardially perfused with 4% paraformaldehyde (PFA) in 0.1 M phosphate buffer (pH 7.3). Brains were removed, post-fixed for 90 min in 4% PFA and cryoprotected overnight in 30% sucrose solution, and 40 μm-thick coronal sections were prepared. For chromogenic immunostaining, sections were quenched for endogenous peroxidase activity by incubation in 3% H_2_O_2_ (Sigma) diluted in methanol for 10 min at 4□°C. Sections were blocked in 10% normal goat serum (Invitrogen), 0.1% Triton-X100 in PBS for 1 h at room temperature. Sections were incubated with primary antibodies for 48 h at 4□°C and biotinylated secondary antibodies (Vector Labs) for 2 h at room temperature. After incubation with ABC reagent (Vector Labs) for 1 h at room temperature and visualization in 3, 3’-diaminobenzidine tetrahydrochloride (DAB; Vector Labs), sections were mounted on Superfrost plus slides (Fisher Scientific), dehydrated with increasing ethanol concentrations and xylene, and coverslipped using Entellan (Merck). All images were captured using a light microscope (Axio Imager M2, Zeiss) equipped with a color CCD camera (AxioCam, Zeiss).

For fluorescence immunostaining, sections were blocked with 10% normal goat serum (Invitrogen), 0.4% BSA and 0.2% Triton-X100 in PBS for 1 h at room temperature. Sections were incubated with primary antibodies for 48 h at 4□°C and secondary antibodies conjugated to AlexaFluor-488, AlexaFluor-594 or AlexaFluor-647 (Life Technologies) for 2 h at room temperature. Sections were mounted on Superfrost plus slides (Fisher Scientific) and coverslipped using Prolong mounting medium containing DAPI (Invitrogen). Images were captured using a Nikon A1plus-RSi laser-scanning confocal microscope (Nikon Instruments) equipped with a 100x oil objective.

### Stereological quantitation of neurons

Unbiased stereological estimation of the number of TH-positive dopaminergic neurons and total Nissl-positive neurons in the substantia nigra pars compacta was performed using the optical fractionator probe of the StereoInvestigator software (MicroBrightField Biosciences), as previously described (30, 32, 38). Every fourth serial coronal section of 40-μm thickness sampled throughout the entire substantia nigra region was immunostained with anti-TH antibody and counterstained with cresyl violet (Nissl) stain. For mice, sampling was performed in a systematic random manner using a grid of 120 × 120 μm squares covering the substantia nigra overlaid on each section and applying an optical dissector consisting of a 50 × 50 × 20 μm square cuboid. For rats, a grid of 220 × 240 μm squares covering the substantia nigra was used. Analyses were performed by investigators blinded to each genotype.

### Quantitative pathology analyses

Digital images of 40 µm-thick sections at 20x magnification were captured using a ScanScope XT slide scanner (Aperio) at a resolution of 0.5 µm/pixel. Image quantitation was conducted manually using the Area Quantification and Microglial Activation modules in HALO analysis software (Indica Labs Inc.) as previously described (45, 49). Analysis thresholds were optimized for each immunohistochemical stain (pSer129-αSyn, GFAP, Iba1 and CD68 in the nigra, or striatal TH) to provide broad detection of pathology in regions with both high and low density pathology, without the inclusion of background staining. Tissue sections spanning the ipsilateral and contralateral substantia nigra or striatum were outlined manually and quantified for the positive staining percentage area occupied by the different immunostains with the positive pixel algorithm, sampled across 2-4 randomly selected images per animal. Positive staining of different immunohistochemically-labeled pathologies was set to a baseline threshold on pixel intensity, such that any pixel at that intensity or greater (i.e., darker) was quantified as a pixel-positive area. The number of positive pixels was normalized per area outline for each section to account for outlined region-to-region area variability. All sections/images were batch analyzed in Halo using the same parameters.

## Acknowledgements

We thank the VAI Pathology and Biorepository, Vivarium, and Optical Imaging Cores for technical assistance. We also thank Dr. Bernard Schneider (Bertarelli Foundation Gene Therapy Platform, EPFL, Switzerland) for providing recombinant AAV2/6-αSyn vector.

## Supplementary Figure Legends

**Figure S1. Heterozygous *VPS35* deletion does not alter the initial spread of** _α_**-Syn pathology in the** _α_**-Syn-PFF model. A**) Schematic illustration of brain injection site. Top and bottom show coronal and sagittal planes, respectively. Red line indicates injection path and red dot indicates injection site. Mice were sacrificed 30 days after unilateral intrastriatal injection of 5 μg mouse α-Syn-PFFs. **B**) Representative images of pS129-α-Syn-positive immunoreactivity indicating equivalent α-Syn pathology/accumulation within the ipsilateral SNpc of age-matched *VPS35*^*WT/WT*^ and *VPS35*^*FLOX/WT*^ mice at 30 days post-injection. The contralateral SNpc lacks pS129-α-Syn-positive pathology, as expected. Scale bar: 100 μm. **C**) Immunofluorescent confocal co-localization of pS129-α-Syn-positive pathology in TH-positive dopaminergic neurons of the ipsilateral SNpc from *VPS35*^*WT/WT*^ and *VPS35*^*FLOX/WT*^ mice at 30 days post-injection of α-Syn-PFFs. pS129-α-Syn accumulates equivalently in the soma of dopaminergic neurons between genotypes. Data are representative of *n* = 3 mice/genotype. Scale bar: 15 μm.

**Figure S2: Quantitative neuropathological analyses**. Midbrain sections were digitized and tissue sections spanning the ipsilateral and contralateral substantia nigra (SN) were outlined manually in HALO software for automated quantitation. Once annotated, algorithms were applied that allowed detection of pathology using thresholding by optical density. Thresholds were individually optimized for each immunohistochemical stain to provide broad detection of pathology in brain regions with both high and low density pathology, without inclusion of background staining. Ipsilateral nigra regions are shown for **(A)** pS129-α-Syn, **(B)** GFAP, **(C)** Iba1, and **(D)** CD68 staining/pathology with or without an analysis overlay (red), which was used to quantify the percent area occupied by pathology/staining.

