## Supplementary figures and images for "VPS35 and α-Synuclein Fail to Interact to Modulate Neurodegeneration in Rodent Models of Parkinson’s Disease"

### Supplemental Figure 1

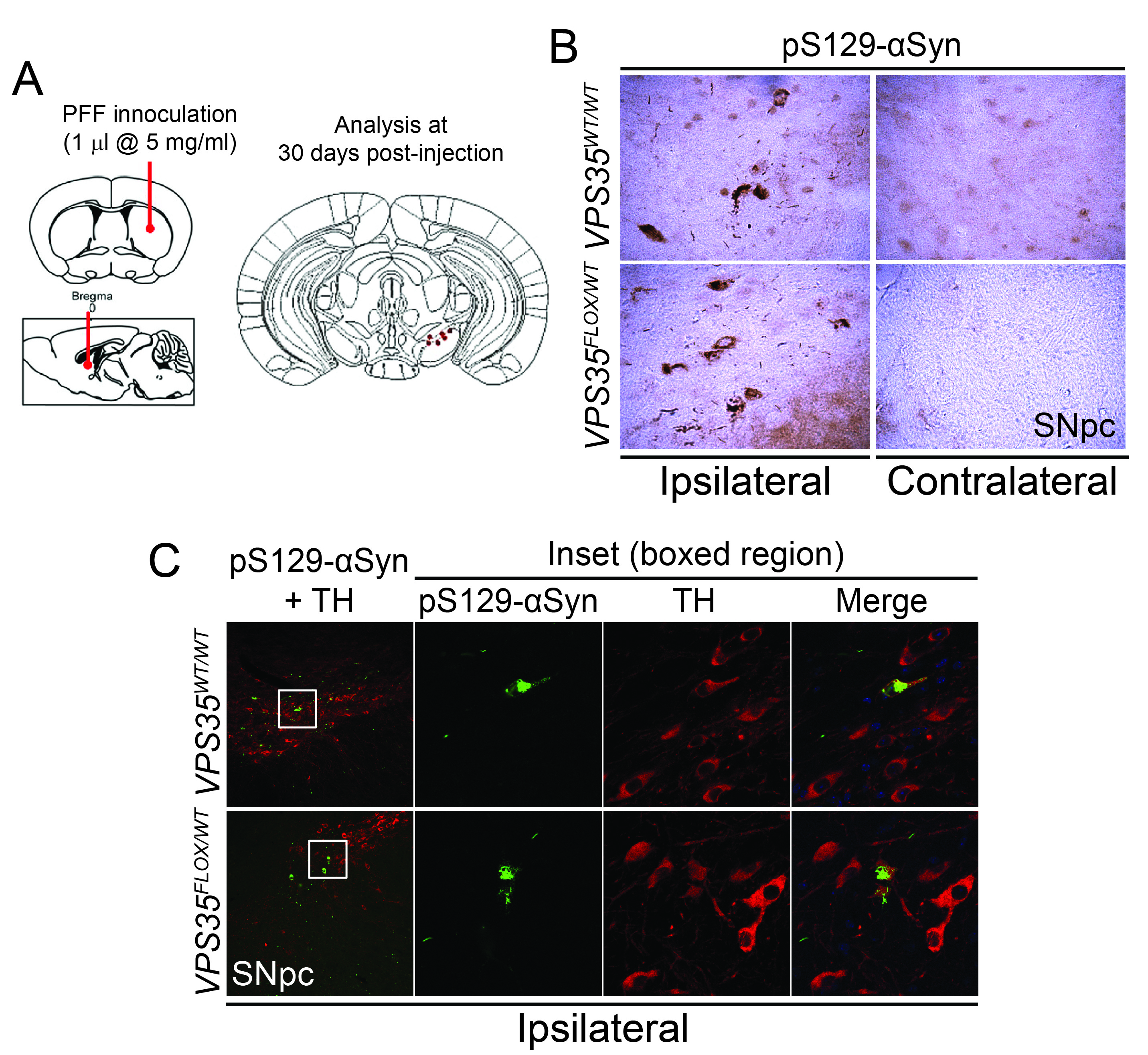

### Supplemental Figure 2

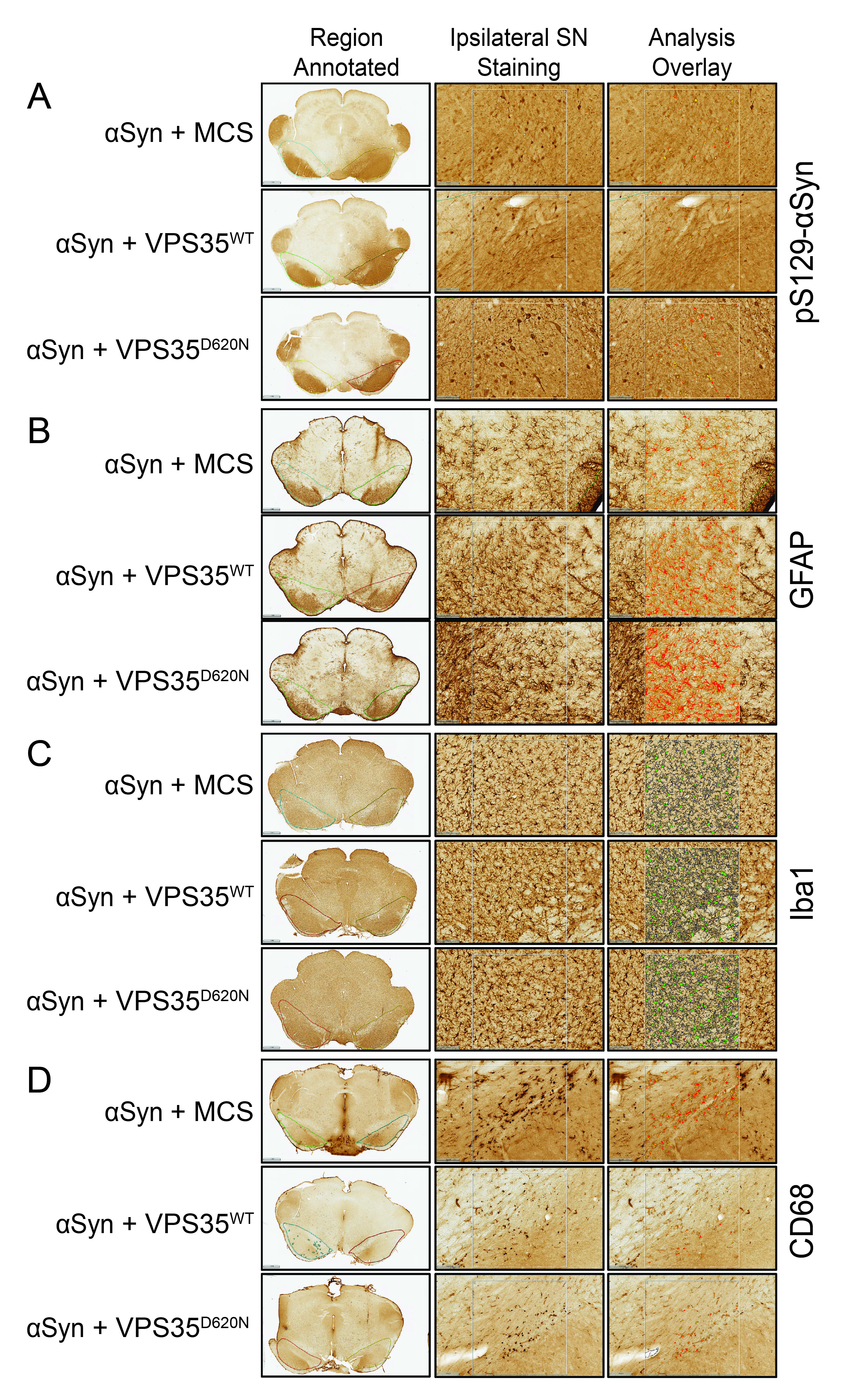
